# Opening a standardized, spatially contiguous biodiversity database collected over 40 years: Czech breeding bird atlases 1973—77, 1985—89, 2001—03, and 2014—17

**DOI:** 10.64898/2026.05.29.728651

**Authors:** Gabriel R. Ortega-Solís, Karel Šťastný, Vladimír Bejček, Tomáš Telenský, Daniela Mellado-Mansilla, Jan Zárybnicky, Florencia Grattarola, Markéta Zárybnická, Zdeněk Vermouzek, Petr Voříšek, François Leroy, Melanie Tietje, Carmen Diana Soria, Elisa Padulosi, Eva Trávníčková, Friederike Johanna Rosa Wölke, Petr Keil

## Abstract

**Motivation:** High-quality biodiversity data with temporal replicates, produced using standardized fieldwork protocols, are rare yet essential for studying long-term biodiversity dynamics. Most available large-scale temporal data only date back one or two decades and/or originate from spatially discrete local observations. Here, we release spatially contiguous, systematically collected, and gridded occurrence data for breeding birds in Czechia, covering the periods 1973—1977, 1985—1989, 2001—2003, and 2014—2017. This database represents the monitoring of ca. 41% of European bird species over 40 years, and it is one of the longest-running nationwide bird-monitoring efforts in the world. We also complement the original data with geospatial metrics to characterize the sampling polygons and provide proxies of sampling effort. By making this dataset openly accessible, we aim to strengthen biodiversity change studies, citizen science, and ornithological research with long-term, highly curated records, backed by well-documented methods, and ready for integration with other datasets.

**Main Types of Variables Contained:** A total of 286302 breeding bird **detections/non-detections** per-grid-cell from 247 species (ca. 41% of the 596 species breeding in Europe). The fourth atlas also contains 9,471 **timed species lists** totaling 276076 additional records collected with standardized effort and partially random spatial sampling on smaller squares dividing the original grid cells.

**Spatial Location and Grain:** Czechia (total area of 78,871 km^2^) covered by a grid of 887 grid cells of 10 by 10 km for the period 1973—77, and 678 cells of 6 minutes latitude and 10 minutes longitude (∼11.2 x 12 kilometers) from 1985 onwards. The timed species lists were collected across 4,851 of 9,844 small squares (∼2.8 x 3 km) that subdivide each original grid-cell into 16 smaller polygons.

**Time Period and Grain:** The sampling years were 1973—1977 (5 breeding seasons), 1985—1989 (5 breeding seasons), 2001—2003 (3 breeding seasons), and 2014—2017 (4 breeding seasons).

**Major Taxa and Level of Measurement:** Birds (Aves). The breeding evidence per species and grid cell was classified following the European Breeding Birds Atlas 2. We provide species-level records matched to the HBW/BirdLife version 9 (2024).

**Format:** The dataset is available for download from Zenodo and is provided as CSV files with fields standardized to Darwin Core, and a GeoPackage file containing all of the spatial grids used. The data are organized into separate files for records and sampling events, corresponding to each atlas. All data are licensed under <u>CC-BY 4.0</u>.

## 1. Background

Biodiversity is facing multiple threats due to global change (IPBES, 2019). Quantifying how biodiversity responds to these threats requires standardized high-quality, large-scale data on temporal trends. However, most standardized biodiversity time series date back only a few decades, and the few existing long-term datasets from older surveys are often not publicly accessible or not even digitized, which hampers the study of long-term trends (Boakes *et al*., 2010). Furthermore, even standardized datasets with long historical coverage cover only spatially disconnected local point observations and transects (e.g. Dornelas *et al*., 2025; Ziolkowski *et al*., 2025), limiting their use for assessing trends at regional and landscape scales.

Among animal taxa, birds are a notable exception in terms of documentation of historical occurrences over large scales, as they are charismatic, conspicuous, and have long attracted biodiversity enthusiasts. This interest drove the establishment of systematic monitoring programmes over continuous spatial grids — known as breeding bird atlases — during the late 1960s and the early 1970s (Udvardy, 1981; Gibbons *et al*., 2007; Dunn & Weston, 2008). These atlases provide a baseline of birds’ richness and species ranges against which newer datasets can be compared. Indeed, some regions have seen re-editions of historical atlases, resulting in unique temporal data (Gillings *et al*., 2019; Keller *et al*., 2020). Most of the repeated atlases only cover 2 sampling periods (editions) (New York State Department of Environmental Conservation, 2000; Keller *et al*., 2020), but there are notable exceptions, such as the 3 editions of bird atlases of the UK (Gillings *et al*., 2019), 3 editions of the Japan atlas (Ueta *et al*., 2021), and 4 editions of the Czech atlas (Šťastný *et al*., 2021) among others. Additionally, some atlases are currently being updated with new editions, such as the New York State, Germany, and the New Zealand atlases.

Breeding bird atlases use standardized survey protocols that are similar across atlas projects, are overseen by expert ornithologists, involve volunteers who are often well-trained in bird identification, and have an expert committee validating the records before they are included in the final dataset (Gillings *et al*., 2019; Keller *et al*., 2020; Šťastný *et al*., 2021). These characteristics make breeding bird atlases suitable for large-scale studies that integrate data from different regions, and particularly attractive for studying species ranges, colonization-extinction dynamics, and broader patterns of biodiversity change. For instance, Thomas & Lennon (1999) compared species ranges between the 1968—1972 and 1988—1991 editions of the breeding birds atlas of Great Britain, and found an average northward expansion of 18.9 km at the northern edge of the ranges, which they linked to climate change. Similarly, Zuckerberg *et al*. (2009) analyzed the 1980—1985 and 2000—2005 atlases of breeding birds of New York, and reported a northward expansion of 11.4 km for species inhabiting the north of the state.

The Czech Breeding Birds Atlas (CzAtlas) is a high-quality, long-term monitoring programme that has so far not been available in a digital, open-access format. It is one of the oldest still-running atlas efforts in the world, and encompasses 4 editions: 1973—1977 (Šťastný *et al*., 1987), 1985—1989 (Šťastný *et al*., 1997), 2001—2003 (Šťastný *et al*., 2009), and 2014—2017 (Šťastný *et al*., 2021). All editions follow a structured fieldwork protocol applied across the entire territory of Czechia (the western part of the former Czechoslovakia). The database also contains counts of volunteers or reporting cards per grid cell as proxies of sampling effort, which can be incorporated into statistical models to account for sampling bias (but see Box 1 for sampling effort caveats). The spatial scope of this atlas overlaps with other large databases, such as the Czech Breeding Birds Survey (Reif *et al*., 2013) and SPARSE (Tschernosterová *et al*., 2023), allowing the use of the Czech birds as a case study to test data integration and interpolation.

In this data paper, we make the four editions of the Czech Breeding Birds atlas, from 1973 to 2017, openly available and comprehensively document them. By providing standardized, spatially contiguous occurrence and breeding-evidence data for 247 species from 153 genera across nearly the entire territory of Czechia, together with detailed information on sampling effort, grid geometry, and taxonomic harmonization, we aim to enable analyses of long-term changes in species’ ranges and community composition at both national and larger scales. Our specific objectives are (i) to describe the historical development and field protocols of the CzAtlas, highlighting changes in sampling design and data sources over time; (ii) to present the structure, content, and standardization of the released database, including its alignment with Darwin Core (and thus GBIF); and (iii) to outline key analytical opportunities and caveats associated with the data, thereby facilitating their integration into macroecological, conservation, and methodological research on biodiversity dynamics.

## 2. Methods

We prepared a cleaned and standardized database, including breeding codes, proxies of sampling effort, and spatial metadata (e.g., measured area and percentage overlap with the country) for every grid cell. The methods for data collection and processing are extensively described in (Šťastný *et al*., 1987, 1997, 2009, 2021); here we provide a shorter consolidated summary.

### 2.1. Study area

Czechia is a landlocked country in the middle of Europe, covering 78,871 km², in the temperate zone of the Northern Hemisphere. The country is at the interface between continental and oceanic climates (Hradecký & Brázdil, 2016). The most relevant topographic features are the mountain ranges along the country’s borders, the central highlands, and the lowlands in the north and southeast (Figure 1). Almost 67% of the territory lies below 500 m.a.s.l., and only 1% exceeds 1000 m.a.s.l. (Statistická ročenka České republiky 2025). The mountain ranges have the lowest temperatures and highest precipitation, averaging less than 3°Celsius and more than 1200 mm, respectively. Around 40% precipitation falls in summer, while in winter it reaches only 15% (Hradecký & Brázdil, 2016). The country’s topography, infrastructure, and population distribution mean the most inaccessible places are in the borderland mountain ranges (Spiekermann *et al*., 2015; Hudeček, 2020).

**Figure 1:**
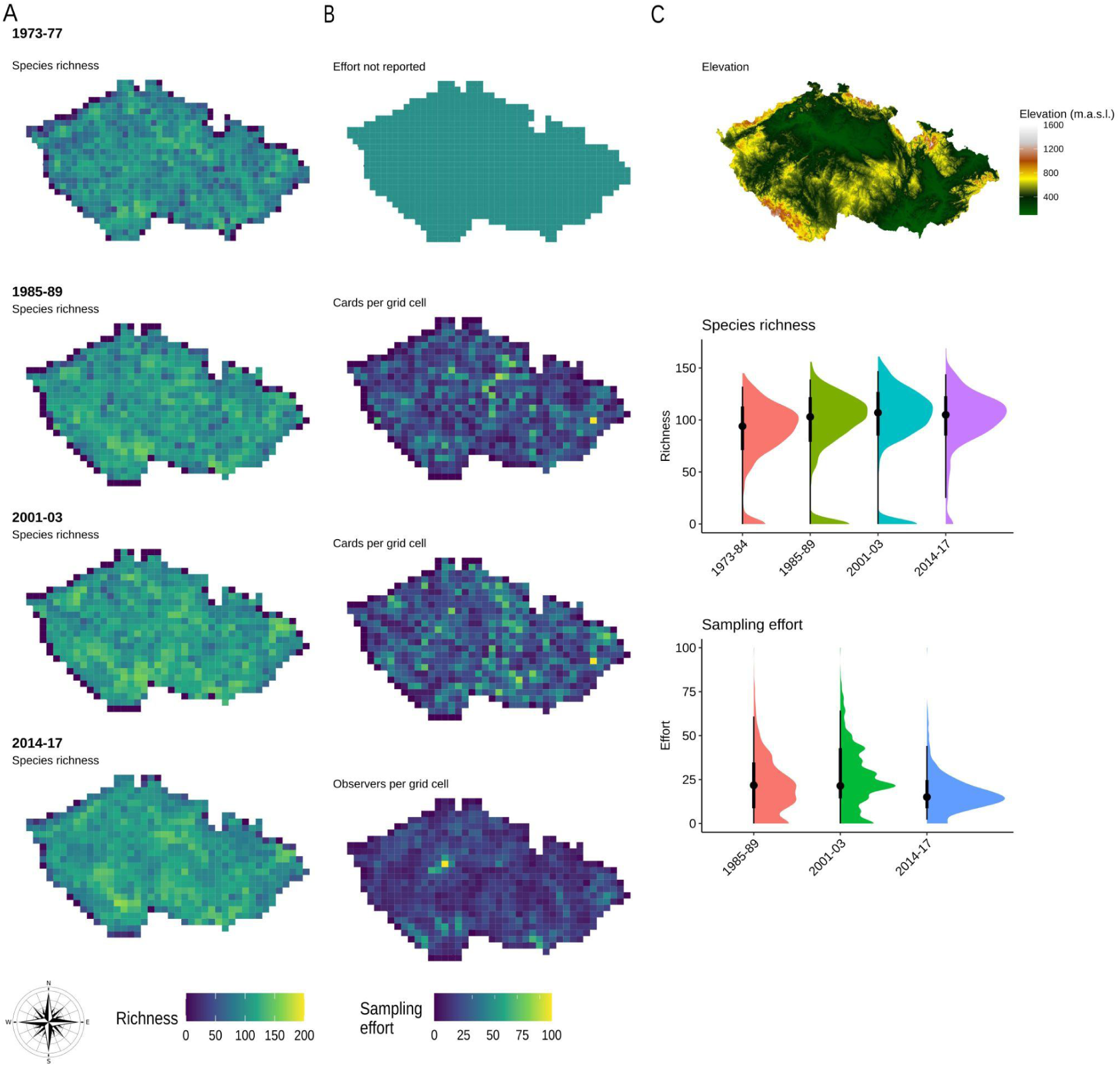
Species richness and sampling effort across the four editions of the Czech bird atlases. Panel A shows the number of species per cell across the atlas editions. B shows the distribution of sampling effort. Effort was not reported during the 1973—1977 sampling period. From 1985 to 2003, the effort was reported as the number of cards per grid cell. In 2014—2017, the effort was reported as the number of observers visiting every cell. Panel C shows the elevation across the country extracted from SRTM digital elevation model (Farr *et al*., 2007), and the distribution of richness and sampling effort per grid cell per atlas edition. Points indicate medians, bars indicate 0.66 and 0.95 intervals. Sampling effort in the bottom-right plot was rescaled to 0—100 for each atlas edition to make them comparable in the plot.

### 2.2. Data collection

Data collection for the first atlas involved approximately 1100 participants, most of whom were members of the Czechoslovak Ornithological Society, joined by anglers, hunters, youth groups, and others. All cells were sampled in the Czech part of the former country during the period from 1973 to 1977, whereas only 51% were sampled in Slovakia. Therefore, this paper focuses exclusively on the Czech data. From 1985 onwards, field surveys were carried out primarily by members of the Czech Society for Ornithology (CSO; www.birdlife.cz/en).

For the first atlas (1973-1977), Czechoslovakia was divided into 1360 grid squares of 10 by 10 km, 887 of which covered the current Czech territory (Figures 1 and S1) (Šťastný et al., 1987). This grid was derived from the Gauss–Krüger kilometre grid in EPSG:4284, without introducing a separate local origin. Individual cells were defined as multiples of 10 km in planar coordinates and coded alphanumerically as a combination of 100 km blocks and their 10 km subdivisions.

From 1985 to 2003, the mapping was based on the system proposed by Ehrendorfer & Hamann (1965). This grid was defined in EPSG:4314, using divisions of 6′ of latitude and 10′ of longitude. It therefore does not represent a rectangular planar grid but rather a system of ellipsoidal trapezoids. For the purposes of faunistic and floristic mapping, this grid was cartographically represented on map sheets at a scale of 1:500,000, with a geodetic basis derived from EPSG:4284.

A new basic mapping grid was introduced during the atlas’s preparation around 2006 with the advent and widespread use of GPS technology. This grid retained the original 6′ × 10′ subdivision but was defined directly in EPSG:4326 (WGS84). It was therefore not a direct transformation of the original grid into EPSG:4326, but rather a redefinition on a different reference ellipsoid and datum. The resulting grid was suitable for working with geographic coordinates in degrees, with grid boundaries aligned to whole minutes of latitude and longitude. However, this approach led to spatial discrepancies relative to the historical grid, locally exceeding 100 m in the field. Given the accuracy of the original mapping, these discrepancies were considered negligible. The rationale for this approach is that a direct transformation between the EPSG:4284 and EPSG:4326 introduces nonlinear shifts and distortions, causing the grid boundaries to no longer coincide with meridians and parallels in EPSG:4326 (Figure S1).

Since 2006, the WGS84-based grid has been used as the primary reference framework. For national cartographic purposes, it is further transformed into the S-JTSK coordinate system (Křovák projection, most commonly in the East North variant, EPSG:5514) using official transformation procedures provided by the Czech Office for Surveying, Mapping and Cadastre (ČÚZK). For atlas purposes, the Czech Republic is divided into 678 grid cells of 6′ × 10′ (Fig. 1) (Šťastný et al., 2021). Sampling covered almost all grid cells, except for a limited number of marginal cells in mountainous border areas that overlap the Czech territory only minimally. The official WGS84-based grid and subdivisions of it can be downloaded from the Nature Conservation Agency of the Czech Republic (https://data.nature.cz/sds/6).

#### Fieldwork protocol

The atlas volunteers were tasked with visiting all habitats within a grid cell, starting with the most dominant and proceeding to the least dominant. Sampling was conducted primarily from dawn to 11 AM on days without rain, fog, or strong winds. Additionally, the observers explicitly searched for rare species in suitable habitats, targeting also crepuscular and nocturnal species. Fieldwork for the first and second atlases lasted five breeding seasons. During the third atlas, the sampling period was shortened to three seasons. Fieldwork for the fourth atlas lasted four breeding seasons. For more details on the fieldwork protocol see Šťastný et al. (2021), pages 13-14.

#### Additional data sources in the fourth atlas

In the fourth atlas (and only there), the field surveys typical for the previous atlases were complemented by two additional data types: (1) Online data sources and (2) timed species lists.

#### Online data sources

Field surveys for the fourth atlas were complemented by *additional records submitted to the faunistic databases* of the Czech Society for Ornithology (2001), the Nature Conservation Agency of the Czech Republic, regional biodiversity monitoring projects (i.e., Breeding Birds Atlas of the Krkonose Mountains), and citizen-science platforms such as eBird, iNaturalist, and Observation.org (Šťastný *et al*., 2021). The additional records included a date, geographic localization, and either a breeding category or detailed breeding notes from which a breeding category could be inferred. Breeding evidence category of those records was discussed with the data providers before inclusion in the atlas database.

#### Timed species lists

The second type of additional data in the fourth atlas are *timed species lists* (TSL), which are lists of species collected with standardized effort and partially random spatial sampling. A TSL contains all species detected within a one-hour walk in a small square in a new grid of small squares (ca. 2.8 x 3 km) created by splitting the basic mapping squares (11.2 x 12 kilometers) into 16 smaller squares (Figure S2), resulting in 9844 small squares. We provide this finer grid in the data repository. Every year, within each mapping square, three of the sixteen small squares were randomly selected for field sampling (new selection was done every year). In some border squares, <3 small squares were selected. Every year, surveyors completed two TSLs in each of the three small squares, one during the early (10th April to mid-May) and one during the late part of the breeding season (mid-May till the end of June) during suitable weather (no strong wind or rain); in the end 8.9 % of TSLs were recorded outside of this range or outside of the prescribed dawn-11am daytime. To maximize the data volume, observers were encouraged to also collect TSLs outside of their officially assigned mapping squares, preferably in the unattended randomly chosen small squares, but any TSLs performed in any small square were also encouraged. The field data collection protocol followed the “timed species counts” of (Pomeroy, 1992; Pomeroy & Dranzoa, 1997): Each 1-hour TSL was split into six 10-minute intervals. The observers took an arbitrary route within the 2.8 x 3 km square, ideally starting at a different place every time, in the habitats most prevalent in the square. Within each TSL, occurrence of all species was recorded, along with the ordinal number (1 to 6) of the first 10-minute interval during which the species was observed. Highest breeding code within the TSL was also recorded. All collected records were checked for errors (typos, inexperience, misunderstanding of methods, etc.) and cleaned from suspicious records by the coordinators. All original records prior to the cleaning are available in the Faunistic database (Czech Society for Ornithology, 2001). In the end, 317 observers collected 9471 TSLs; out of these, 67% from randomly sampled squares. Out of the 9844 small squares, 4851 (49 %) were covered by TSLs. Out of these, 74 % were covered by random sampling. In the majority of cases, one or two TSLs per small square were performed.

#### Breeding evidence categories

The breeding evidence in the database follows the European Breeding Birds Atlas 2 (EBBA2) classification, using the categories 0 (not nesting), A (possibly nesting), B (probably nesting), and C (nesting), along with breeding behavior codes ranging from 0 to 16 (see descriptions in Table 1). Atlases 1, 2, and 3 used a different coding system, with categories ranging from A (not breeding) to D (confirmed breeding), which were subsequently harmonized to the EBBA2 scheme (see Table 1).

**Table 1:** Breeding codes used in every atlas. The fourth atlas used categories standardized to the European Breeding Birds Atlas 2.

| Category<br>standardized<br>to EBBA2 | Category<br>atlases 1,2,<br>and 3 | Category<br>definition | Code | Behaviour description |
| --- | --- | --- | --- | --- |
|  | A | Non-breeding | 0 | Species observed during the surveys, but suspected to be still on migration or not to breed during the surveyed period |
| A | B | Possibly breeding | 1 | Species observed in the breeding season in a suitable nesting habitat (but some waders and gulls may stay in such habitats through the season without breeding, so further evidence is needed). |
| A | B | Possibly breeding | 2 | Singing male(s) or other vocalizations indicating breeding, heard in the breeding season. |
| B | C | Probably breeding | 3 | A pair (male and female) was observed in a suitable nesting habitat in the breeding season. |
| B | C | Probably breeding | 4 | Permanent territory inferred from territorial behaviour (display, song, chasing rivals, etc.) recorded at the same site on at least two occasions a week or more apart. |
| B | C | Probably breeding | 5 | Courtship display, mate-attraction behaviour, or copulation was observed. |
| B | C | Probably breeding | 6 | Bird seen prospecting or searching for possible nest sites. |
| B | C | Probably breeding | 7 | Agitated behaviour or alarm-calling of adults suggesting the presence of a nest or young nearby. |
| B | C | Probably breeding | 8 | Presence of a brood patch on captured adult(s). |
| B | C | Probably breeding | 9 | Adults seen nest-building or excavating a nesting hole. |
| C | D | Confirmed breeding | 10 | Distraction displays, such as feigning injury to draw attention away from the nest or young. |
| C | D | Confirmed breeding | 11 | Used nest (occupied or obviously recently occupied) or fragments of eggshells found. |
| C | D | Confirmed breeding | 12 | Recently fledged young (for species that feed young) or down-clad young (for precocial species) were found. |
| C | D | Confirmed breeding | 13 | Adults seen entering or leaving a presumed nest site under circumstances indicating an occupied nest (including high or hidden nests or cavities) or adults apparently incubating. |
| C | D | Confirmed breeding | 14 | Adults carrying faecal sacs away from a nest or bringing food to young. |
| C | D | Confirmed breeding | 15 | Nest with eggs found. |
| C | D | Confirmed breeding | 16 | Nest with young found (seen or clearly heard). |

#### Records validation

After data collection, the records in each atlas edition were double-checked and validated by atlas coordinators and experts for specific bird species groups. The timed-species-lists (fourth atlas) also includes records of species not breeding (breeding code 0) that we excluded from the main dataset. If a species was seen breeding in a grid cell at least in one season, it was recorded as breeding for the whole period of the corresponding atlas.

### 2.3. Auxiliary data

Here, we also provide auxiliary data that are not part of the original four atlases, but users will likely still appreciate having them at a single repository with the original data. We matched the species names and taxonomic information to HBW/BirdLife 9 (HBW and BirdLife International, 2024), and extracted the primary habitat and diet for each species from Birdbase 2025.1 (Şekercioğlu *et al*., 2025). We used a recent phylogenetic tree (McTavish *et al*., 2025) to explore the phylogenetic representativeness of the CzAtlas compared to the European Breeding Birds Atlas (Keller *et al*., 2020). Finally, we also calculated the area, side length, and North-South and East-West extension of each cell before and after cropping it with the Czech national borders using PostgreSQL/PostGIS (PostgreSQL Global Development Group, 2024; PostGIS Project Steering Committee, 2025).

### 2.4. Database structure

Every record in our database corresponds to species-cell-sampling period, that is, *“an occurrence of a species, within a grid cell, during a given atlas sampling period”.* Species in this case correspond to the original identification used in each atlas edition, including subspecies. For example, if *Corvus corone* was observed in a cell during courtship and the subspecies *Corvus corone corone* was also found nesting in that cell, the species appears twice in the database.

We standardized variable naming according to the Darwin Core standard when an equivalent name existed, using the R package galaxias (Westgate *et al*., 2026). We transformed the spatial grids into EPSG:4326 coordinate reference system with terra 1.8 (Hijmans, 2026). All the information is provided in CSV files per atlas period, split into species records (records_*_cleaned.csv files) and sampling event (events_*.csv files) related metadata. The spatial grids are stored in a single GeoPackage (grids.gpkg). All data management and standardization procedures were done in R 4.5 (R Core Team, 2025).

**Table 2:**
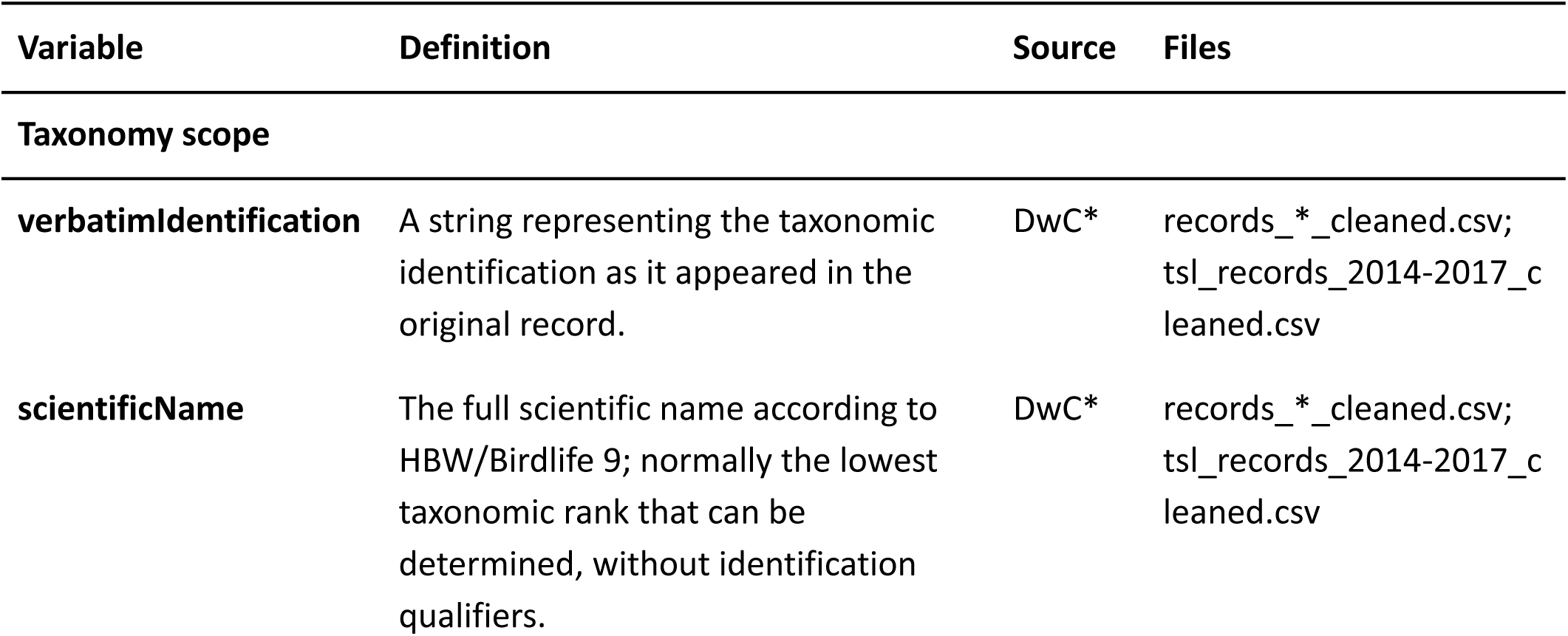
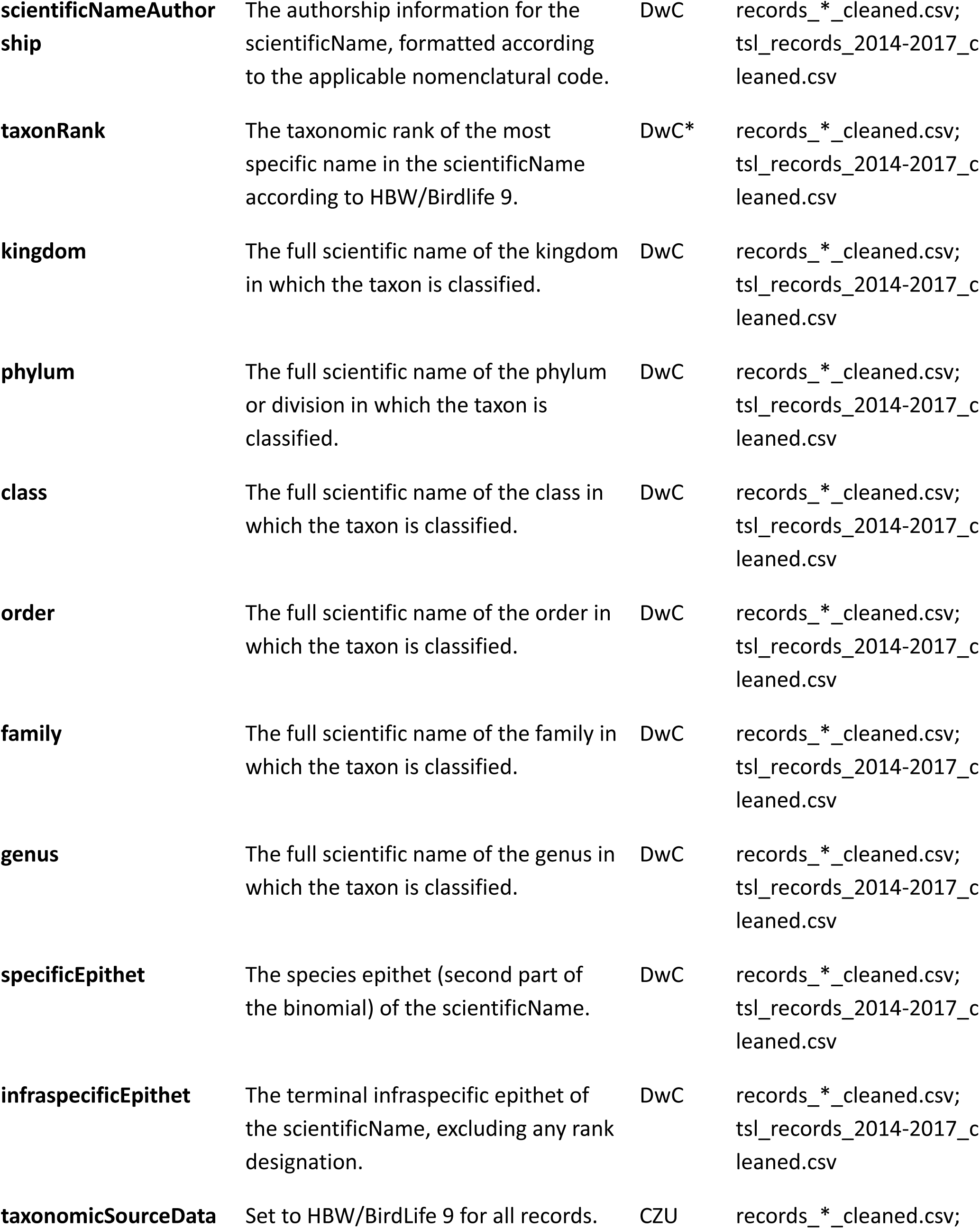
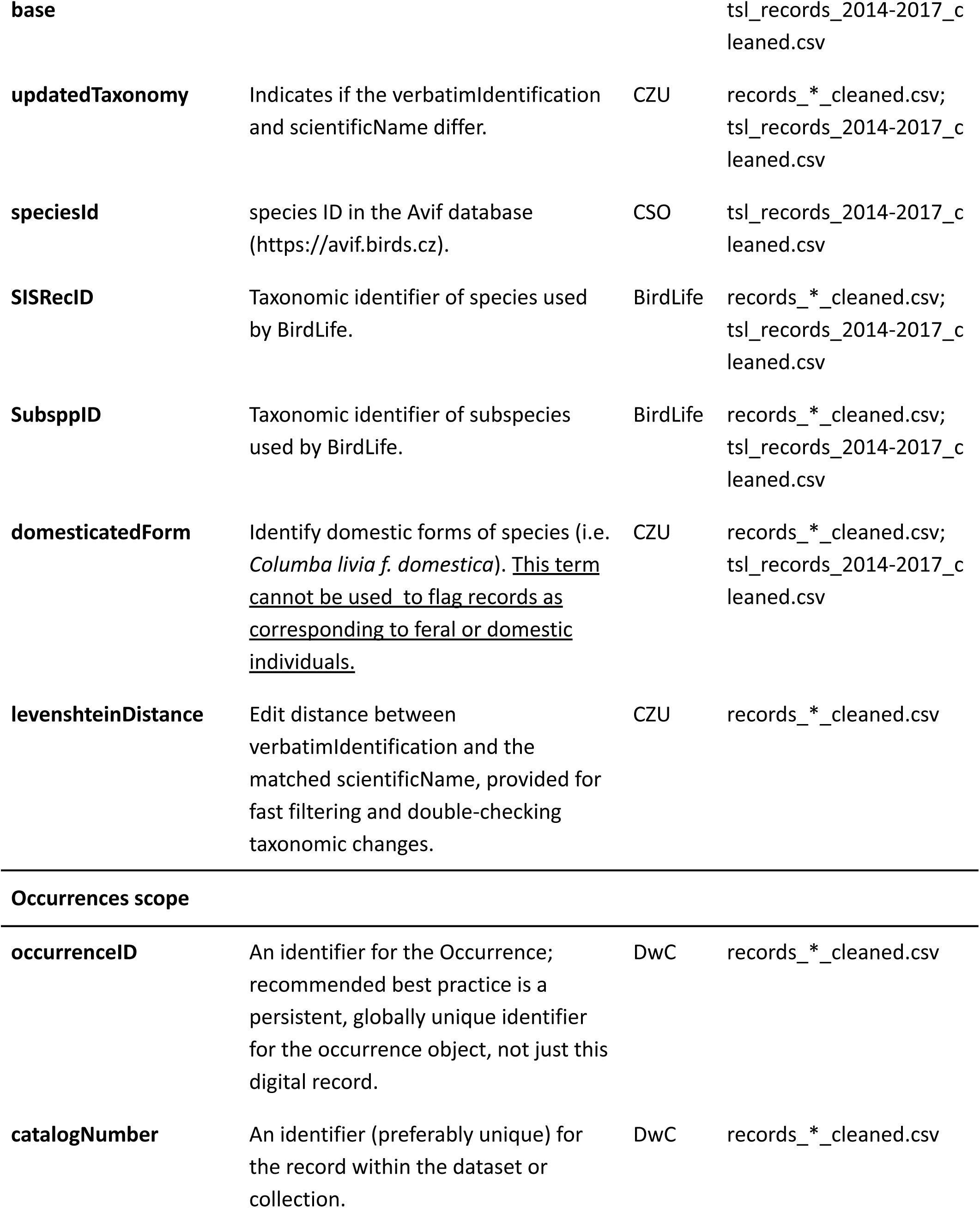
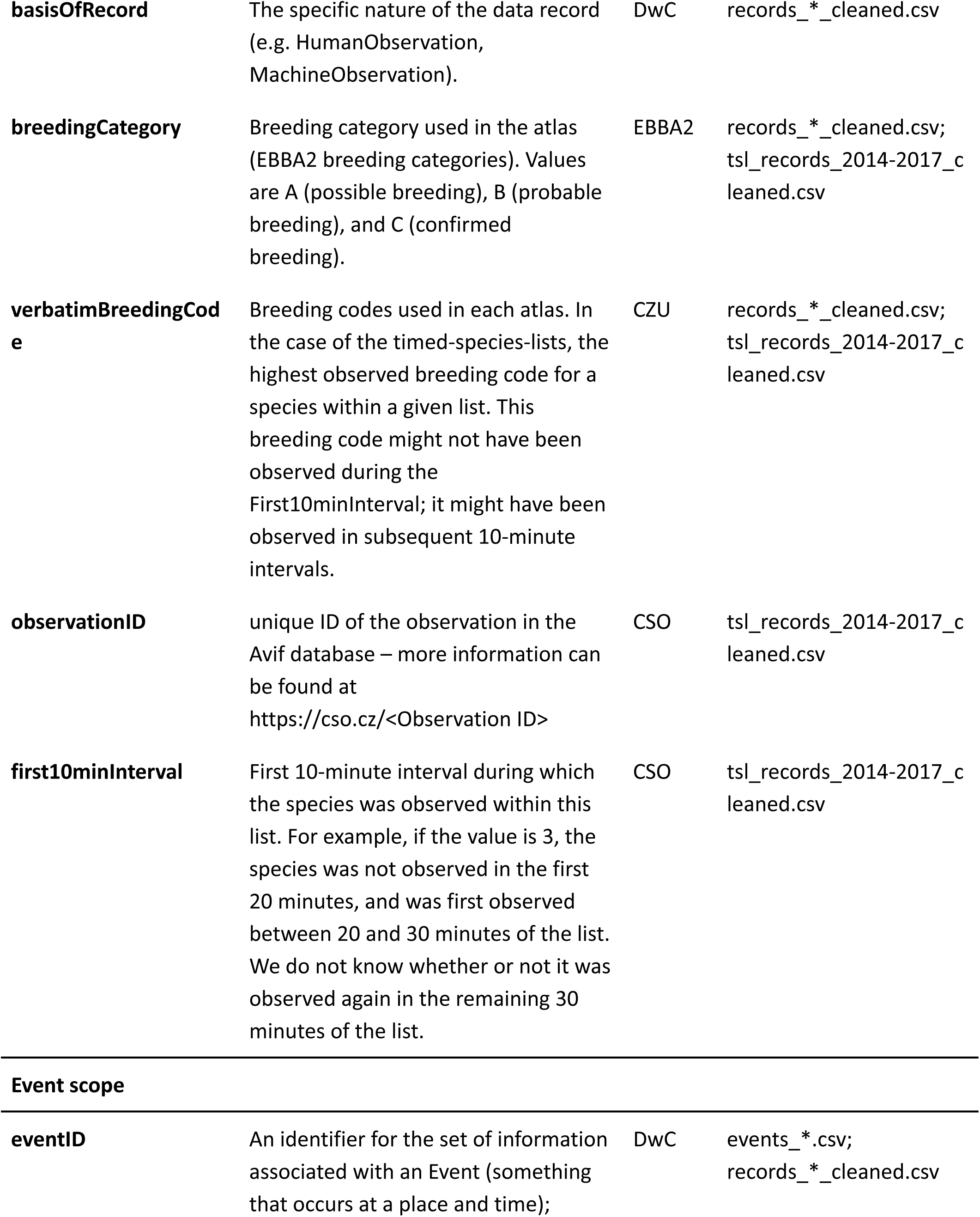
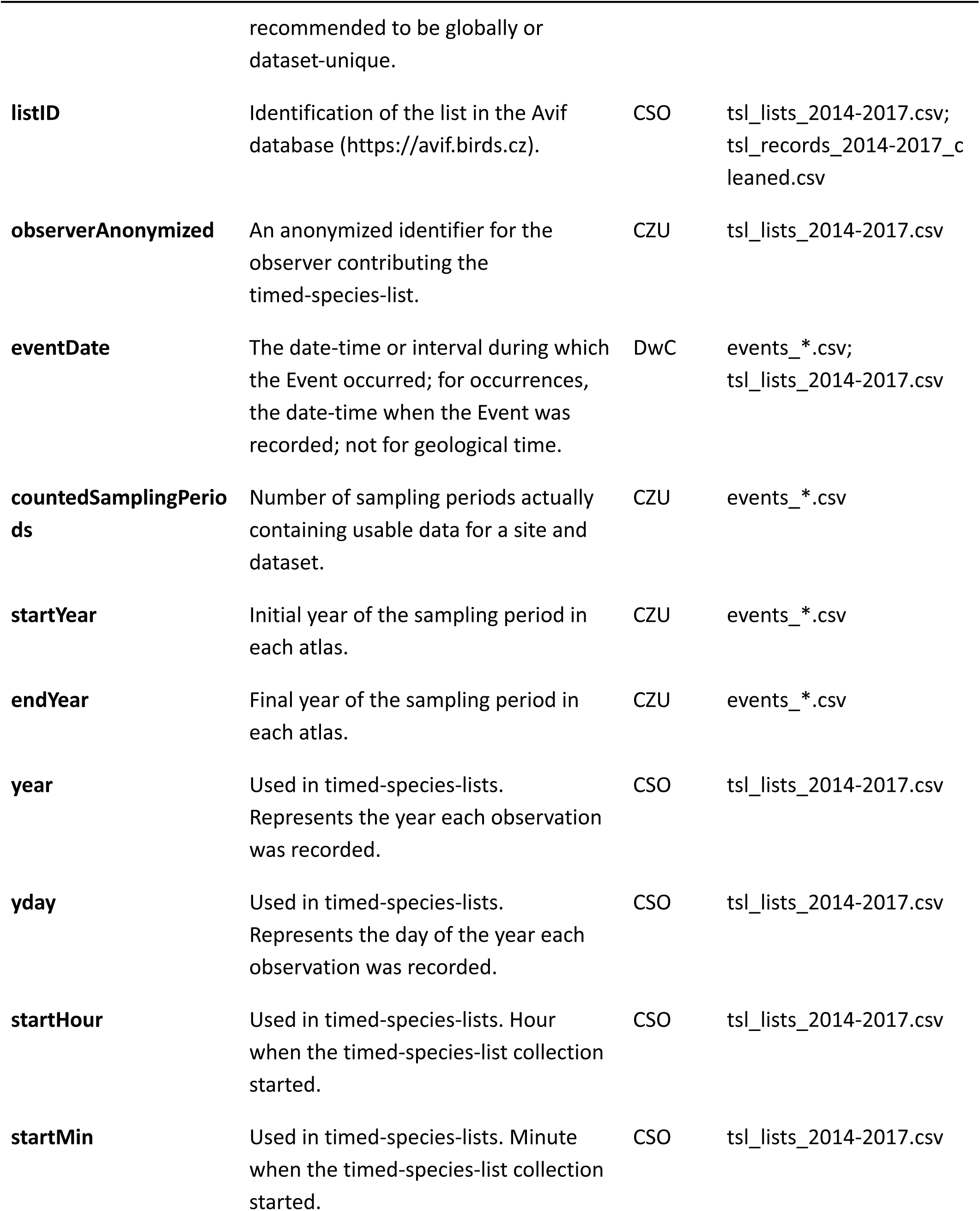
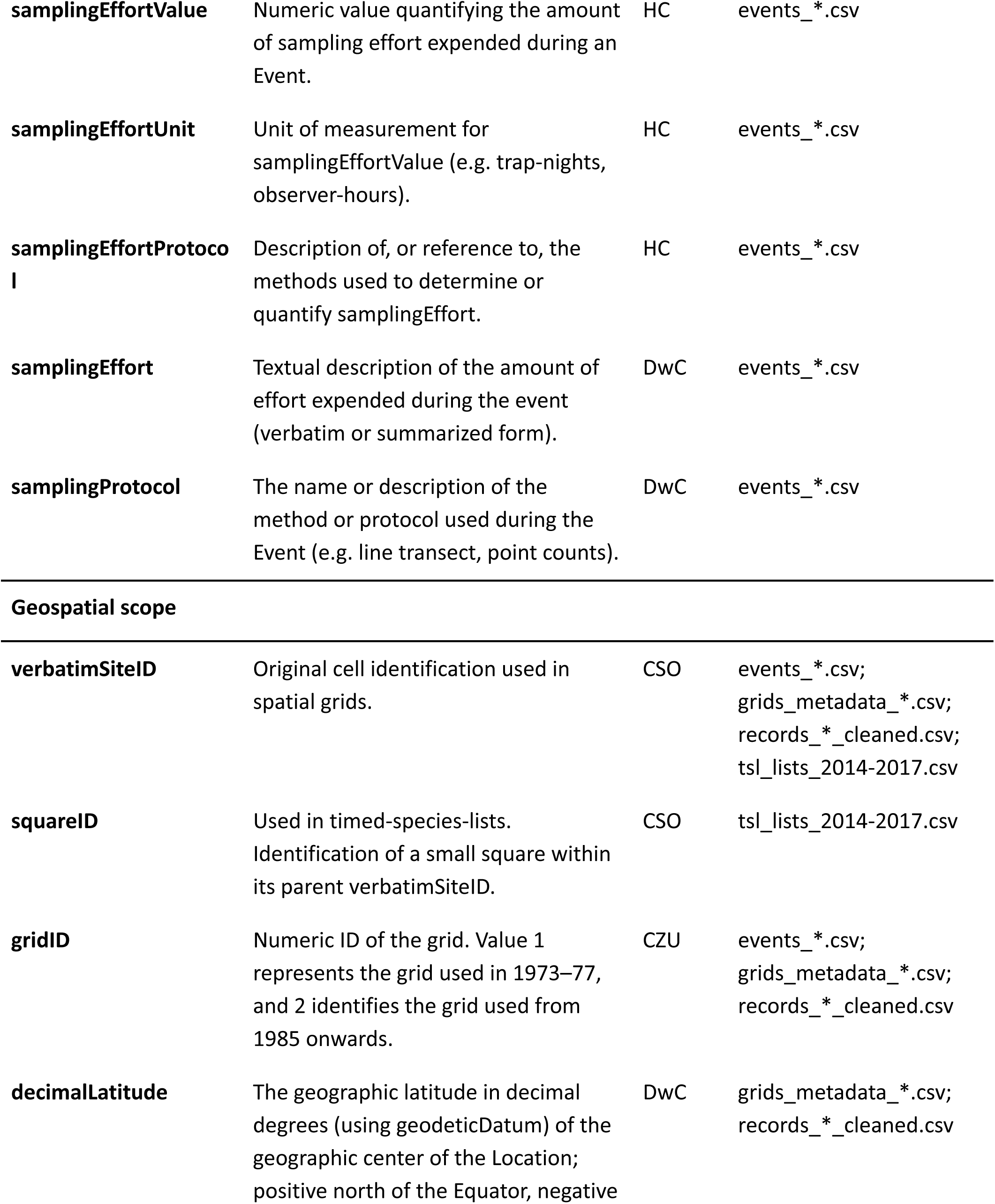
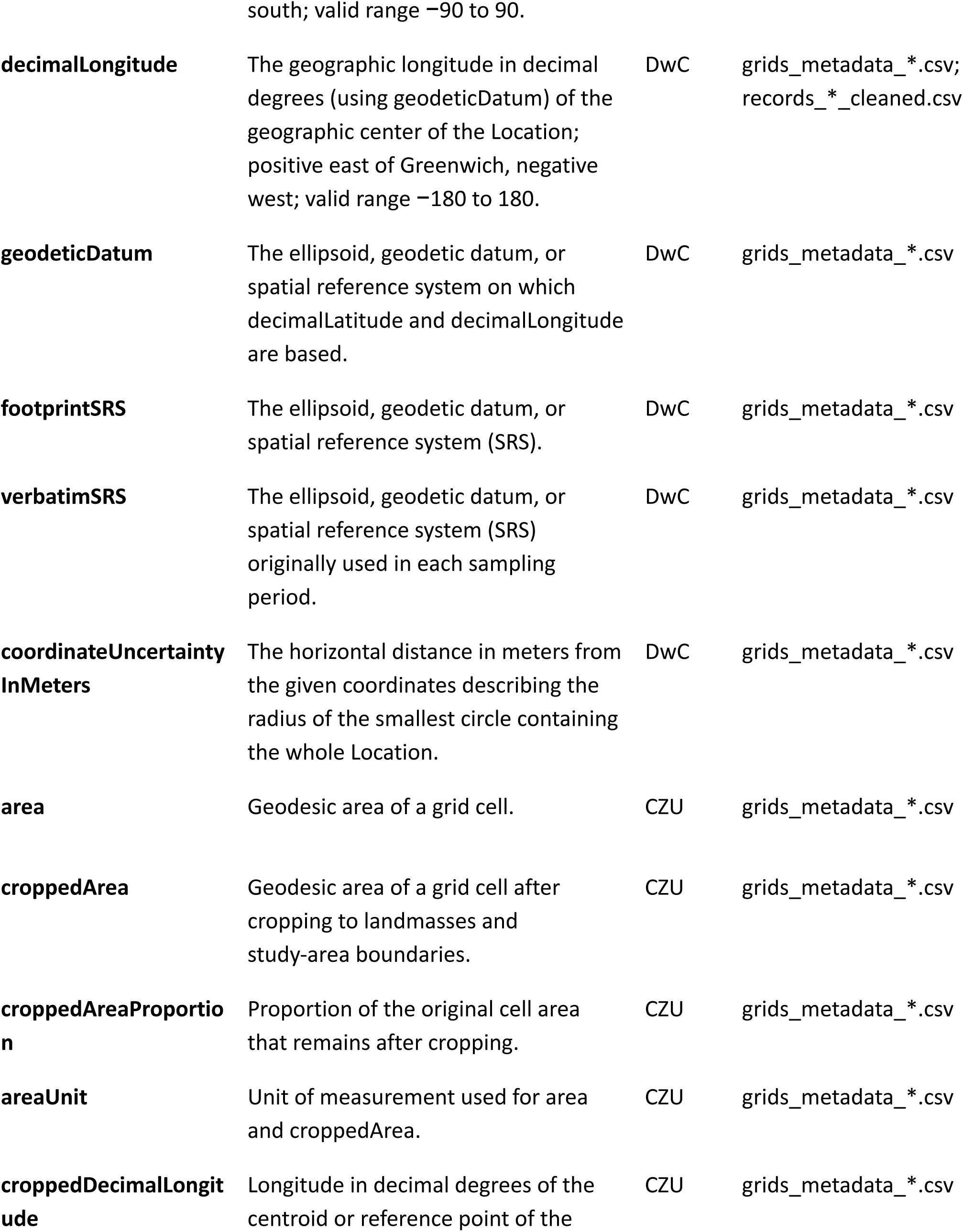
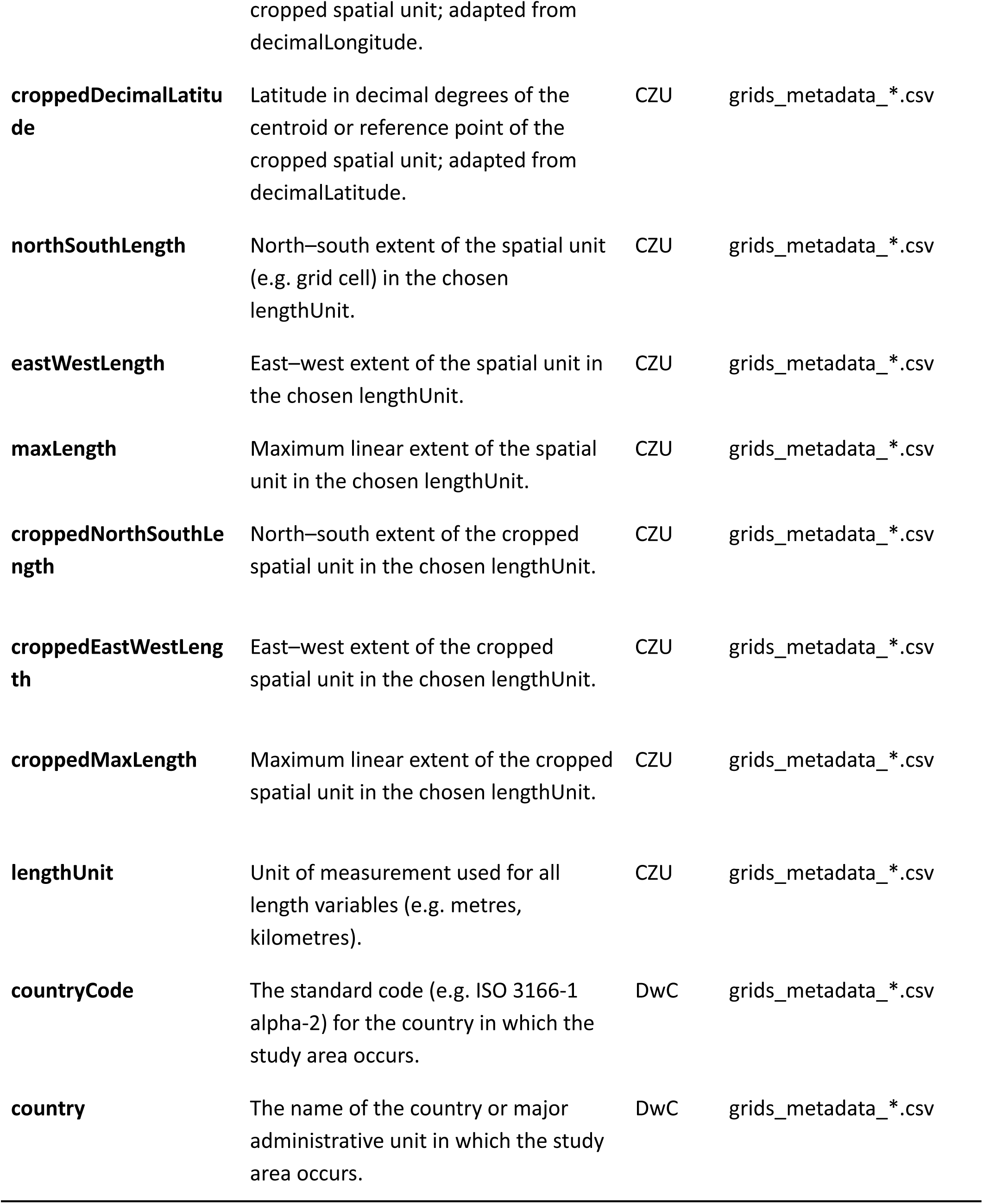
Variables in the database. Source shows the origin of the variable names and definitions used. Sources were Darwin Core (DwC, Darwin Core Maintenance Group, 2021), Humboldt Core (HC, Guralnick *et al*., 2018), Birdbase 2025.1 (Şekercioğlu *et al*., 2025), BirdLife (BirdLife International, 2026), Czech Society for Ornithology (CSO), and EBBA2 (Keller *et al*., 2020); CZU shows variables defined at the Czech University of Life Sciences. Variables adapted from other sources are identified by a star (*). The column “Files” indicates the files that contain each variable.

## 3. Data overview

### 3.1. Spatial and temporal distribution of records

The spatial distribution of per-grid-cell species richness across the four sampling periods is stable (Figure 1). Within each sampling period, effort per grid cell is distributed more or less randomly across the country, without clear spatial aggregation patterns, but with a tendency to decrease in remote areas near the country borders (Figure 1). The fourth atlas includes records from previously unsampled border cells.

### 3.2. Taxonomic, phylogenetic, and ecological representation

The database comprises 247 species spanning 153 genera, 57 families, and 21 orders. Contrasted with the 596 breeding bird species found in Europe (Keller *et al*., 2020), Czechia accounts for 81% of the orders, 67% of the families, 52% of the genera, and 41% of the species on the continent (Figure 2). Passeriformes constitute 43% of species in Czechia, followed by Charadriiformes (11%), Anseriformes (10%), and Accipitriformes (6%), while all the remaining orders reach less than 5% (Figure 2). Czech birds are distributed across the entire phylogenetic tree of European birds (Figure 2), representing all major lineages. Most species are wetland inhabitants (28%), followed by forest and woodland species (22% and 17% respectively). Regarding trophic levels, roughly 68% are carnivores (grouping invertivores, vertivores, piscivores, and scavengers), 17% consume plants, fruits, seeds or nectar, and 15% are omnivores (see for percentages per order Figure S3).

**Figure 2:**
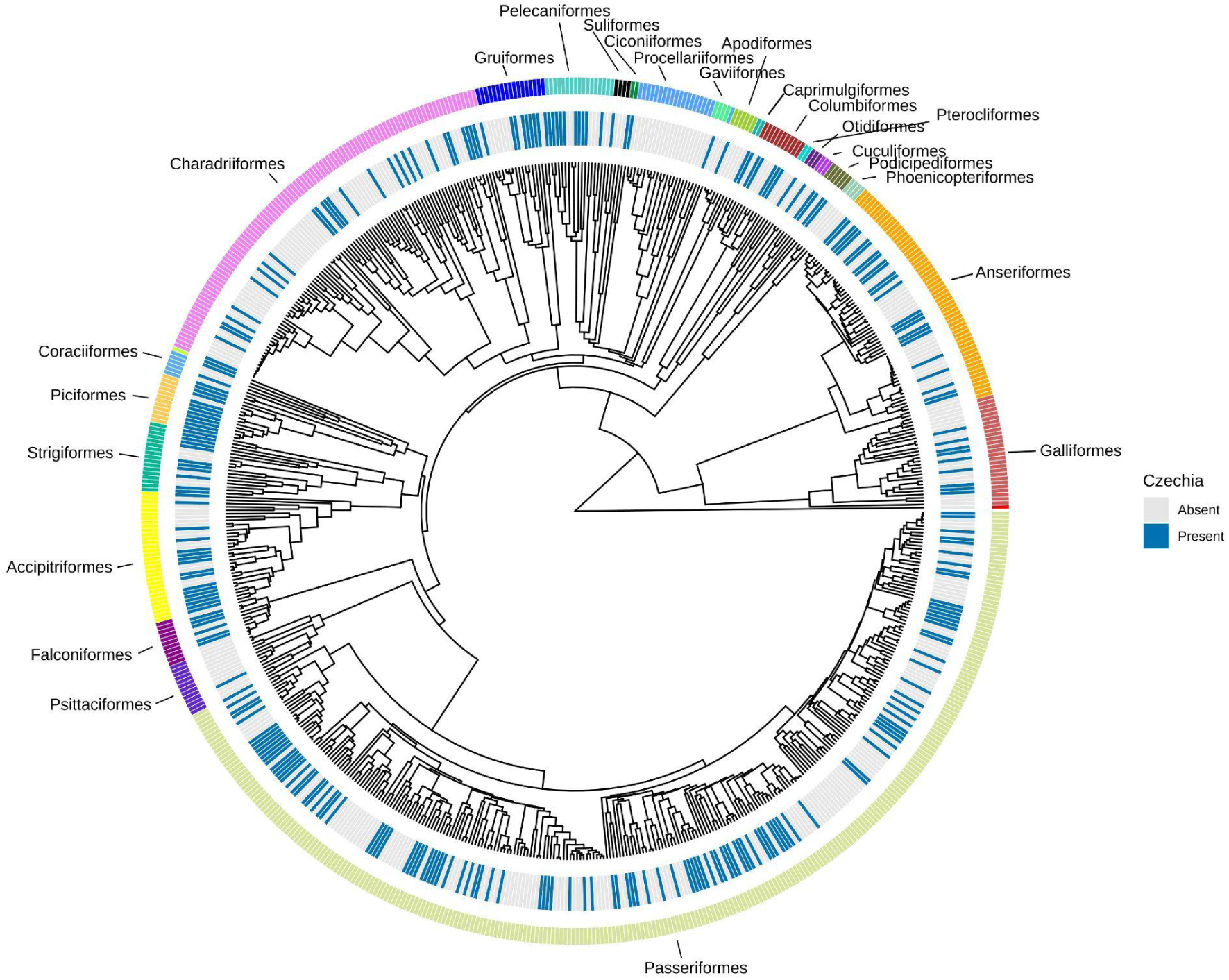
Taxonomic distribution of bird species occurring in Czechia. The phylogenetic tree of (McTavish *et al*., 2025) was pruned to include only bird species recorded in Europe (Keller et al., 2020). The inner ring highlights species present (cyan) and absent (grey) in Czechia. The outer ring and accompanying labels denote the taxonomic orders of birds.

## 4. Usage and Future Notes

### 4.1. Past research and future opportunities

The Czech bird atlas data have been extensively used in macroecological research. Several studies have used only one edition of the atlas. For example, Storch & Šizling (2002) used the data to show that the core-satellite hypothesis applies at large scales. (Storch *et al*., 2003) used them to test mechanisms driving the species-area relationship. The data were used to develop new approaches for species distribution modelling (Moudrý & Šímová, 2013; Moudrý *et al*., 2017; Gábor *et al*., 2022), and to test hypotheses about drivers of biodiversity (Prajzlerová *et al*., 2024). There were also case studies of individual species’ ecology based on the data (e.g.(Ševčík *et al*., 2021)).

More recently, researchers have begun leveraging the temporal dimension of the data. This includes analyses of simple trends of species occupancy (Reif *et al*., 2013). Keil et al. (2018) then used it to demonstrate the scale-dependence of extinction rates. Using the last three editions of the data, Leroy *et al*. (2024) showed that colonization rates increase towards coarser scales, while extinction rates peak at intermediate grains, producing the strongest richness gains at coarse grains. Soria *et al*. (2026) found that, while the spatial autocorrelation of species richness and distributions did not change over time, its strength decreased with increasing grain size. The latest study using multiple editions of the atlases focused on temporal scaling of ecological patterns and their drivers (Mellado-Mansilla *et al*., 2026). All these studies relied on the spatial contiguity and repeated coverage of the atlas’s data.

Importantly, the four temporal replicates in the CzAtlas offer opportunities for new research. Future research can address long-term range shifts, changes in species richness, temporal turnover, and colonization and extinction dynamics. The standardized grid facilitates comparisons with other atlases and national monitoring schemes. Information on the breeding status of multiple species should enable analyses of temporal shifts in breeding behavior. Thanks to the high phylogenetic and functional representation of the CzAtlas, the data can be used to test hypotheses about evolutionary and functional mechanisms. As in the above-mentioned studies (Keil *et al*., 2018; Leroy *et al*., 2024; Soria *et al*., 2026), spatially contiguous grids can be used to study phenomena that spatially discrete data (e.g. BioTime; (Dornelas *et al*., 2025)) cannot capture. Examples are spatial configurations of species distributions and diversity, such as their spatial fragmentation and aggregation (clumping, autocorrelation), fractality, or elongation. The timed species lists in the fourth atlas also enable fine-grained species distribution modeling at ∼3km small grid cells and serve as a complement to the main atlas data.

### 4.2. Caveats

#### Variation in temporal span and sampling effort

The temporal span of each atlas period ranged from three breeding seasons (2001—2003) to five seasons (1973–1977 and 1985–1989), with the fourth atlas spanning four seasons (2014—2017). This variation should be taken into account. For instance, studying trends in a species’ reproductive status may depend on identifying the maximum possible breeding evidence, which in turn can be related to the number of breeding seasons in each atlas. Further, the incorporation of supplementary data sources in the fourth atlas likely increased detection probability relative to earlier periods by adding opportunistic records to the standardized sampling. However, the database provides useful proxies of sampling effort—number of reporting cards per grid cell for the second and third atlases, and number of observers per cell for the fourth atlas—which can be incorporated as covariates in statistical models to account for effort-related biases. We must stress that researchers using this dataset must be aware that effort proxies are not perfect and may still have methodological limitations. For instance, a highly skilled observer may record more species in a cell than many less experienced observers visiting another cell. In addition, a high number of observers does not always mean meaningfully higher effort, because popular sites may attract many birdwatchers who repeatedly report the same species. The TSL are available only for the fourth atlas sampling period (2014—2017). Also note that 8.9% of TSL data fall outside the proper time window of the survey protocol; therefore, applying date/time filters that differ from those described in the **Timed species list** section may affect comparability with the model outputs presented in the fourth atlas.

#### Change in the spatial grid

The change in grid projection between the first and subsequent atlases (Figure S1) can pose a challenge, as it precludes direct cell-to-cell comparisons. For multi-period analyses requiring spatially consistent units, users can consider three general options: (i) exclude the first atlas from the temporal series, retaining the three later periods on a common grid; (ii) analyze each atlas independently; or (iii) apply spatial interpolation or resampling procedures to project the first atlas onto the later grid.

#### Border cells

Grid cells along international borders may overlap only partially with Czech territory, resulting in effective survey areas that are substantially smaller than the nominal cell size. To quantify this, we provide a “croppedCellArea” field in the GeoPackage, which specifies the area of each cell within the national boundary. Users are encouraged to evaluate whether to include border cells in their analyses—particularly those with very small overlap—given that reduced area may confound richness and occupancy estimates. The database also includes the cell side lengths and the north–south and east–west extension of each cell, both before and after cropping, to facilitate informed decisions.

### 4.3. Contact and collaboration

For better interpretation and use of the data, users are encouraged to contact the data custodians (K. Šťastný, V. Bejček, or the Czech Society for Ornithology) or their associates prior to analysis. The raw data from the fourth atlas, which were collected electronically via Avif, are preserved in the Avif database (https://avif.birds.cz). Detailing which species and periods are being analysed will facilitate dialogue and help prevent misinterpretation.

## DATA AVAILABILITY STATEMENT

The data accompanying this article are openly available in Zenodo at https://doi.org/10.5281/zenodo.18503904. The code is available on the GitHub repository https://anonymous.4open.science/r/Czech_birds_data_paper-B8E3/.

## ACKNOWLEDGMENTS

This study stands on the shoulders of the thousands of volunteers who participated in the CzAtlas. PK, GOS, DMM, CS, MT, FW, FG, and EP were funded by the European Union (ERC, BEAST, 101044740). Views and opinions expressed are, however, those of the author(s) only and do not necessarily reflect those of the European Union or the European Research Council Executive Agency. Neither the European Union nor the granting authority can be held responsible for them.

We dedicate this work to the 100th Anniversary of the Czech Society for Ornithology (2026) and to all the ornithologists working in Czechia during the last 100 years.

## BIOSKETCH

Karel Šťastný and Vladimír Bejček are both professors of zoology and ornithology, and are the main coordinators behind the four editions of the atlases (together with the deceased Karel Hudec). The Modelling of Biodiversity Lab (MOBI Lab) is a research group focused on exploring the temporal and spatial dynamics of biodiversity and how they are affected by human-driven environmental change. The team aims to identify the patterns and mechanisms that shape the geographic distribution and temporal changes of biodiversity using large-scale ecological data.

