## Supplemental tables and figures for "Opening a standardized, spatially contiguous biodiversity database collected over 40 years: Czech breeding bird atlases 1973—77, 1985—89, 2001—03, and 2014—17"

**SUPPORTING INFORMATION**

**Table S1:** Summary of species and records per atlas.

| Atlas Period | Grid Type | Cell Area | N Cells | N Species | Total Records | Survey Years |
|---|---|---|---|---|---|---|
| 1973-1977 | 10×10 km | ~100 km² | 887 | 193 | 80023 | 5 |
| 1985-1989 | 6'×10' | ~134 km² | 678 | 206 | 65598 | 5 |
| 2001-2003 | 6'×10' | ~134 km² | 678 | 213 | 68849 | 3 |
| 2014-2017 | 6'×10' | ~134 km² | 678 | 242 | 71832 | 4 |
| | | | Total | 247 | 286302 | |

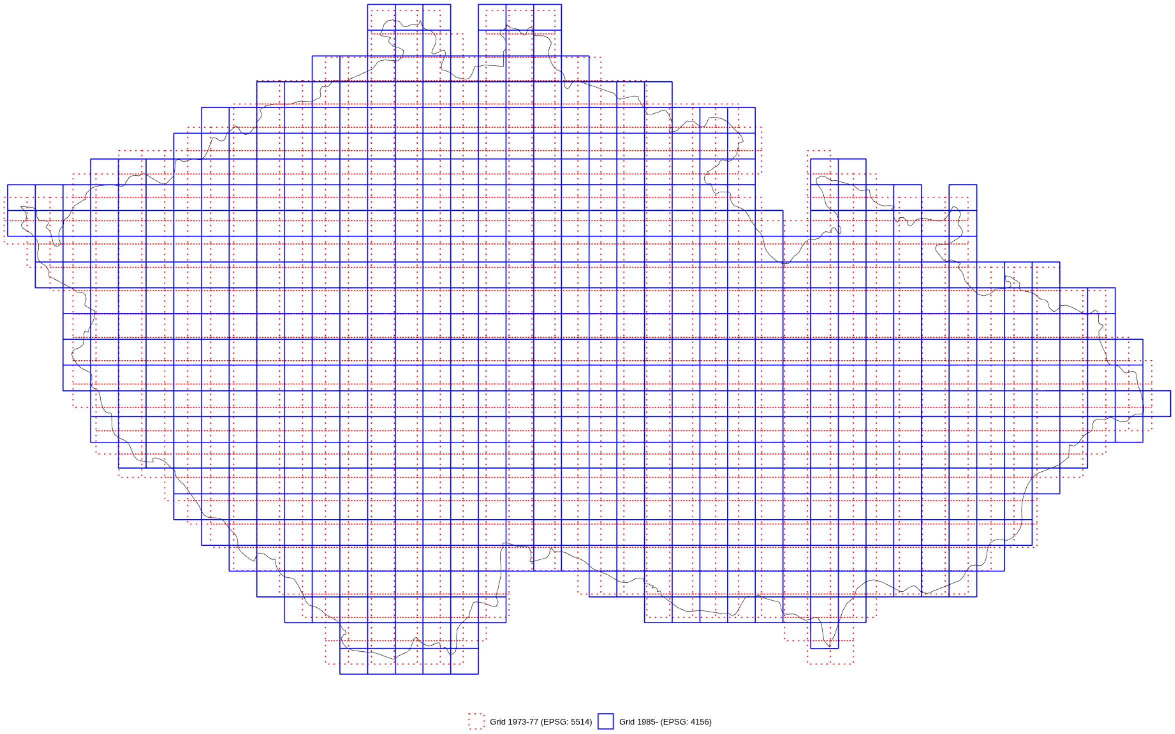


**Figure S1:** Spatial grids used for the first (1973-77) and newer (1985 onwards) atlases.

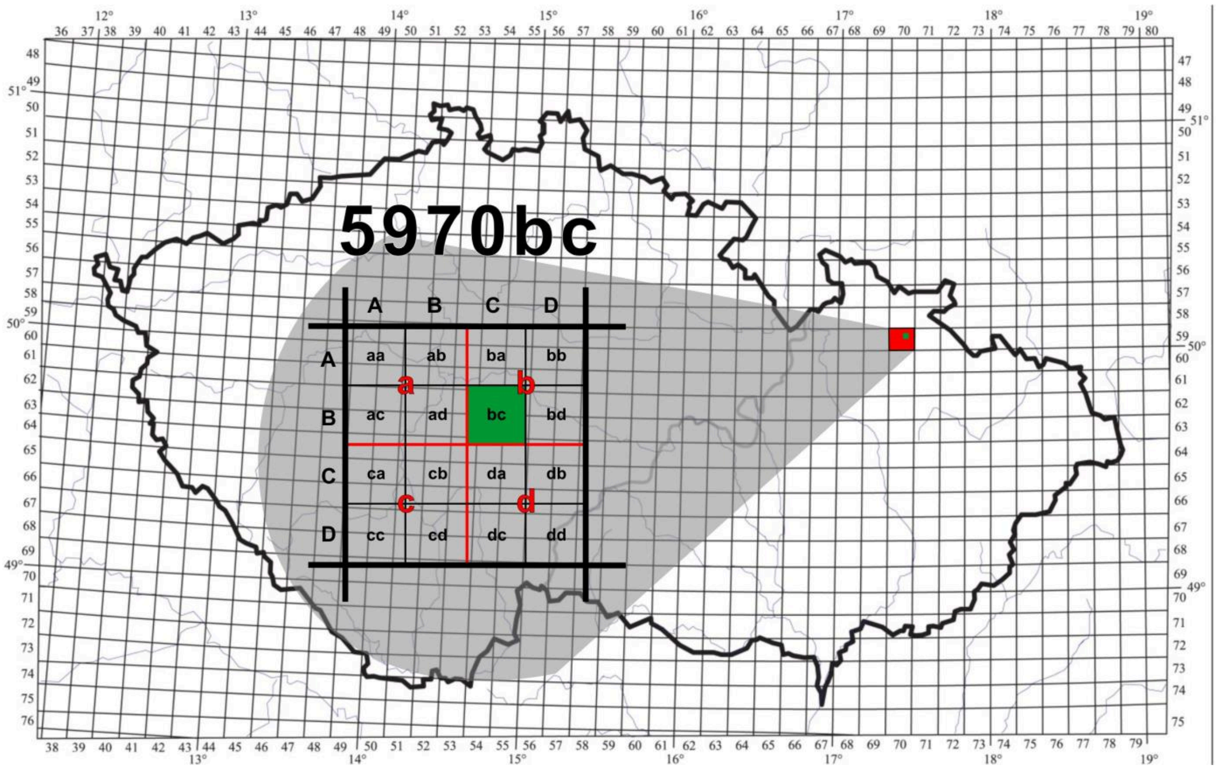


**Figure S2**: A schematic map showing the division of a basic mapping square into a grid of 16 small squares.

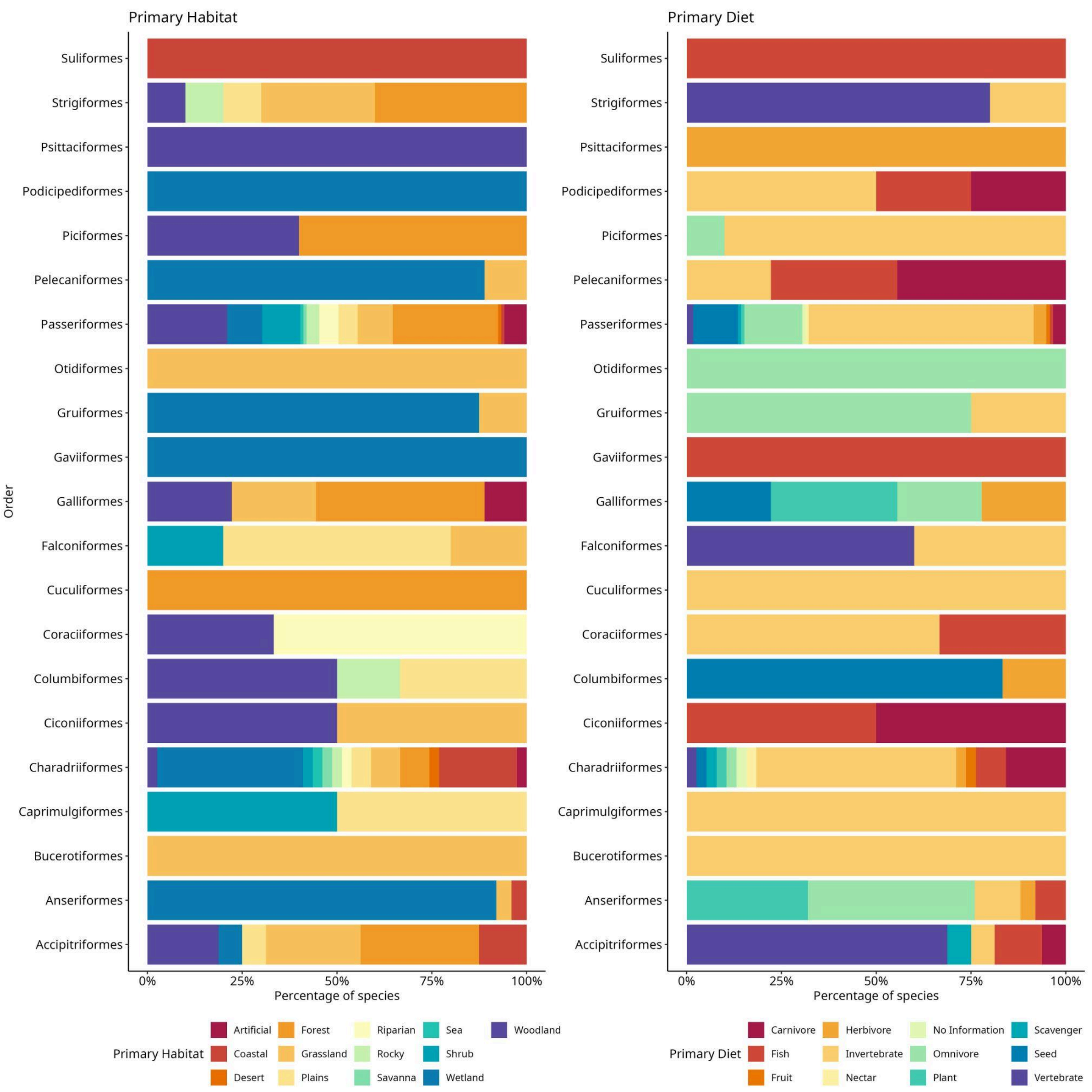


**Figure S3:** Percentages of primary habitat and diet categories per order. Categories follow (Şekercioğlu et al., 2025).
